# MONTE enables unified pan-cancer tumor purity estimation and methylation correction from bulk DNA methylation arrays

**DOI:** 10.64898/2026.01.22.701164

**Authors:** Mirae Kim, Wei-Hao Lee, Vicky Yao

**Author notes:** These authors contributed equally to this work (Mirae Kim, Wei-Hao Lee).

## Abstract

Bulk DNA methylation profiling is widely used to study cancer epigenomics in clinical settings, but these measurements aggregate signals from malignant and non-malignant cells, introducing composition-dependent confounding that complicates tumor-intrinsic interpretation and cross-cohort analyses. While existing methods can estimate tumor purity and, in some cases, correct methylation measurements, they typically require cancer-specific reference models, matched normal samples, or predefined probe sets, limiting their applicability to rare cancers, different clinical cohorts, and cross-dataset comparisons. We present MONTE (Methylation-based Observation Normalization and Tumor purity Estimation), a unified, cancer label-free framework for tumor purity inference and CpG-resolved methylation correction from bulk DNA methylation data. MONTE learns probe-wise relationships between methylation and tumor purity using an empirical Bayes-moderated linear model and infers purity in new samples via signal-to-noise weighted aggregation, without requiring matched normals, cancer labels, or predefined probe sets. A single pan-cancer MONTE model outperforms existing cancer-specific methods for purity estimation across 21 cancer types, generalizes across purity references, and runs orders of magnitude faster on full-dataset analyses. MONTE also introduces Bayesian transfer learning, which enables efficient recalibration to alternative purity definitions, validated on three independent external cohorts. Methylation correction with MONTE further amplifies tumor-relevant regulatory signal and improves the reproducibility of differential methylation analyses. By unifying purity estimation and correction in a single flexible, scalable, and interpretable framework, MONTE broadens the accessibility of tumor-intrinsic methylation analysis across cancer types and datasets.

## Introduction

DNA methylation is a major epigenomic modification that plays a central role in regulating gene expression and maintaining cellular identity. These patterns are dynamically shaped by both exogenous environmental exposures, such as smoking, and endogenous biological factors, such as age. In cancer, aberrant methylation landscapes are a hallmark of tumorigenesis, often involving the silencing of tumor suppressor genes and activation of oncogenic pathways, thereby promoting uncontrolled cell proliferation and tumor progression [1]. Interpreting these alterations in bulk tumor samples, however, is complicated by the fact that measured methylation profiles reflect mixtures of malignant and non-malignant cell populations (e.g., immune and stromal cells) [2].

Most cancer DNA methylation studies rely on bulk tumor samples, in which measurements aggregate signals across heterogeneous cellular compositions. DNA methylation microarrays remain widely used in both research and clinical settings due to their robustness and cost-effectiveness. These platforms quantify methylation at hundreds of thousands of predefined genomic sites using probes that target cytosine-guanine (CpG) dinucleotides. However, variation in tumor purity can introduce systematic differences in the observed methylation profiles. This compositional heterogeneity can bias downstream analyses, leading to spurious or attenuated differential methylation signals and confounded clinical associations [3–5]. Accurate tumor purity estimation followed by appropriate correction of methylation measurements is a critical prerequisite for analyses that aim to recover tumor-intrinsic epigenetic signals.

Several approaches have been proposed to infer tumor purity from DNA methylation data, but each is constrained by some combination of three requirements: matched normal samples, cancer-specific models, or predefined probe sets (Table 1). InfiniumPurify [5], MEpurity [8], and PAMES [7] rely on differential methylation between tumor and tissue-matched normal references to identify cancer-associated CpG sites, then estimate purity using summary statistics [5, 7] or mixture modeling [8] over these predefined, cancer-specific probe sets. While effective in supported settings, these methods require appropriate normal samples and are trained separately for each cancer type, limiting their applicability in rare cancers and in cohorts lacking suitable normal references. Regression-based approaches relax some of these requirements but introduce others. RF_purify [9] applies random forest regression directly to bulk methylation but requires an exact match to its training probe set, limiting robustness to probe filtering and cross-dataset application. PureBeta [6] does not require predefined probe sets, but it combines probe-wise linear regression with a computationally intensive mixture modeling procedure and requires cancer-specific model training. Pretrained PureBeta models are currently available for only a small number of cancer types.

**Table 1.** Comparison of MONTE with existing, executable DNA methylation tumor purity estimation methods.

| Property | Method |  |  |  |
| --- | --- | --- | --- | --- |
|  | MONTE | PureBeta [6] | InfiniumPurify [5] | PAMES [7] |
| **Core capabilities** |  |  |  |  |
| Purity estimation | ✓ | ✓ | ✓ | ✓ |
| Methylation correction | ✓ | ✓ | ✓ | ✗ |
| **Data and annotation requirements** |  |  |  |  |
| Does not require normal samples | ✓ | ✓ | ✗ | ✗ |
| Does not require cancer labels | ✓ | ✗ | ✗ | ✗ |
| Does not require predefined probe sets | ✓ | ✓ | ✗ | ✗ |
| **Statistical modeling properties** |  |  |  |  |
| Modeling framework | OLS + EB | OLS | DMC | DMC |
| Probe-weighting by statistical reliability | SNR | N/A | N/A | N/A |
| **Transferability and usability** |  |  |  |  |
| Transfer learning across datasets/purity definitions | ✓ | ✗ | ✗ | ✗ |
| Software availability | Python | R | R | R |

Note: MEpurify and RF\_purify are not listed because we could not obtain results from them (see Methods). DMC: Differential methylated CpGs; EB: Empirical Bayes; OLS: Ordinary least squares; SNR: Signal-to-noise ratio

Importantly, tumor purity estimation alone can identify samples with substantial non-tumor contamination but cannot recover tumor-intrinsic DNA methylation signals. To make effective use of samples with varying purity, purity information must be coupled with explicit correction of methylation measurements to yield a CpG-resolved pure-tumor methylome, required for downstream analyses such as molecular subtyping [2, 10], regulatory interpretation [11], or integration with other omics layers [12, 13]. Only PureBeta [6] and InfiniumPurify [5] currently support both estimation and correction, but both rely on fixed, predefined reference models that are tightly coupled to specific cancer types and data contexts. These constraints limit adaptability across datasets and purity scoring frameworks, motivating the need for more flexible and scalable approaches.

Here, we present MONTE (Methylation-based Observation Normalization and Tumor purity Estimation), a unified, cancer label-free framework that addresses these limitations by jointly enabling tumor purity estimation and CpG-resolved methylation correction (Figure 1). MONTE models probe-wise relationships between observed methylation and tumor purity using a regression framework with empirical Bayes variance moderation, resulting in stabilized and uncertainty-aware estimates across heterogeneous datasets. Purity inference for new samples is performed via signal-to-noise weighted aggregation of probe-level effects, enabling robust prediction even when only a subset of probes are available, without requiring cancer labels, matched normal samples, or predefined probe panels. To extend beyond a single cohort or purity metric, MONTE incorporates Bayesian transfer learning, allowing pretrained models to be efficiently recalibrated to new datasets or alternative purity scoring systems using limited labeled data. The learned probe-purity relationships further enable projection to complete tumor purity, producing CpG-resolved tumor-intrinsic methylation profiles suitable for downstream analysis. Because no single purity reference is definitive, we evaluate MONTE against multiple reference measures and on independent external cohorts. Across these benchmarks, MONTE outperforms existing cancer-specific methods in purity estimation across 21 cancer types, generalizes across purity references, and produces methylation profiles that improve recovery of tumor-intrinsic signals.

**Figure 1.**
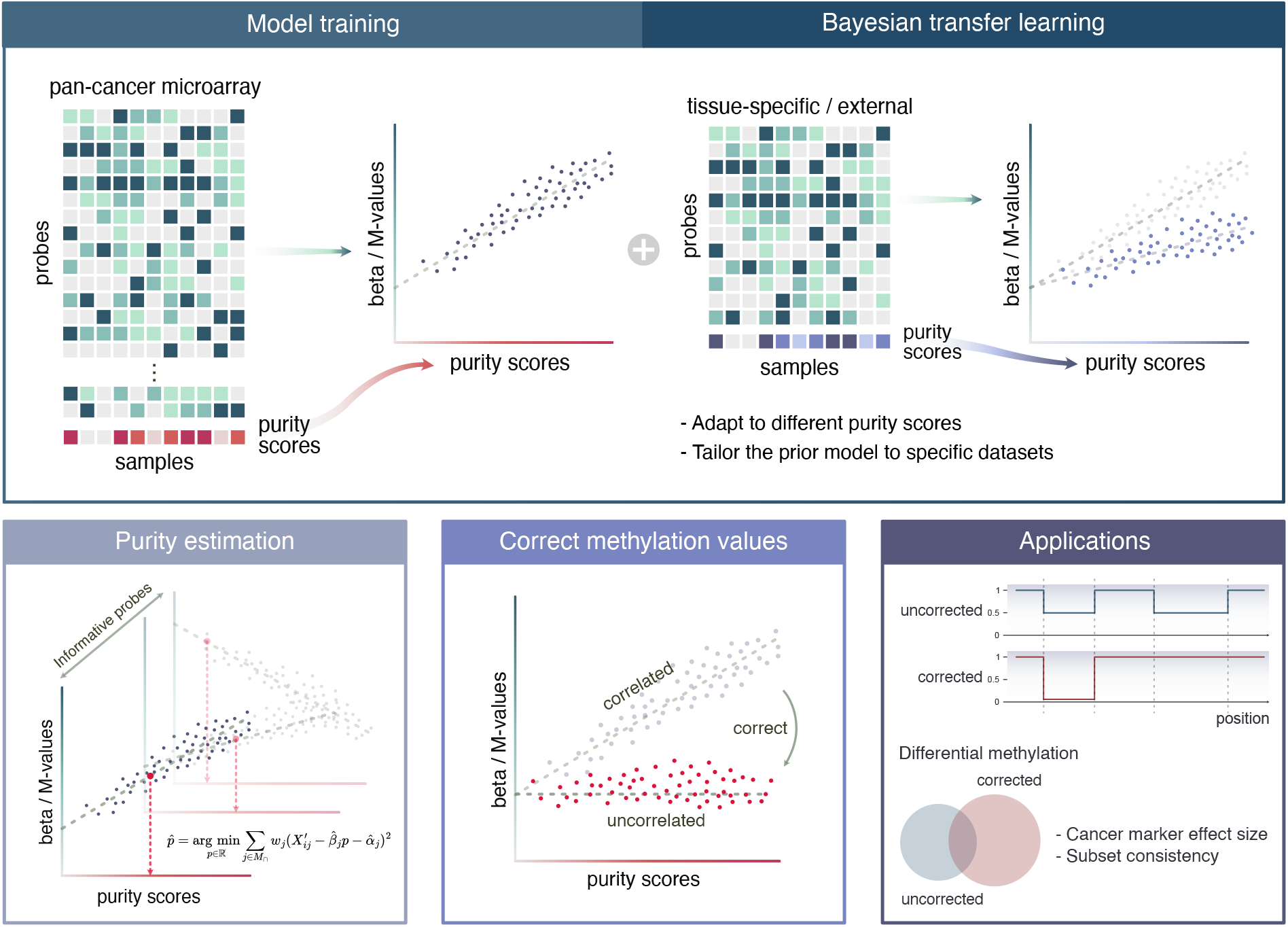
Overview of the MONTE framework. (Top) The pan-cancer model learns probe-purity relationships from bulk DNA methylation data without the use of cancer labels, normal samples, or predefined probe sets. Learned coefficients can be optionally refined through Bayesian transfer learning on tissue-specific, cancer-specific, or external datasets, allowing the model to adapt to alternative purity definitions or new data cohorts. **(Bottom)** MONTE estimates sample-level tumor purity using signal-to-noise weighted aggregation of probe-level effects and corrects methylation values by removing purity-associated signal. The resulting profiles better reflect tumor-intrinsic methylation to improve downstream analyses.

## Results

### A single MONTE model provides robust pan-cancer tumor purity estimation

MONTE estimates tumor purity from bulk DNA methylation data by aggregating probe-level associations between methylation and purity learned in a pan-cancer setting (Figure 1). In this framework, probe-wise regression coefficients are estimated using ordinary least squares with empirical Bayes variance moderation, and probes are weighted by their signal-to-noise ratio when inferring purity for new samples. A top-N probe selection step is used to regularize estimation when large or heterogeneous probe sets are available (see Methods).

We evaluated MONTE using DNA methylation data from The Cancer Genome Atlas (TCGA) [14], which provides one of the largest publicly available DNA methylation resources spanning diverse cancer types.

We used Illumina HumanMethylation 450K data from TCGA samples with consensus purity estimates (CPE), which integrate multiple purity measures derived from copy number (ABSOLUTE), gene expression (ESTIMATE), DNA methylation (LUMP), and histopathology [2]. After quality control, this resulted in a dataset of 7,040 samples across 21 cancer types. CPE provides the most comprehensive purity reference available for TCGA and serves as our primary training target. In addition to ABSOLUTE and ESTIMATE, the respective copy-number-based and expression-based components of CPE that contain no methylation signal, we also evaluate MONTE against whole-genome sequencing-derived purity on external PCAWG cohorts (below). To mitigate concerns about circularity and data leakage, samples were split into disjoint training and test sets using a 70%/30% split stratified by cancer type, and all evaluations were performed exclusively on held-out test samples, notably without providing any cancer labels.

To understand how individual components of the MONTE framework contribute to robust purity estimation, we performed an ablation analysis comparing alternative formulations of the same modeling framework, including baseline ordinary least squares (OLS) regression, empirical Bayes-moderated OLS (Var), empirical Bayes with signal-to-noise (SNR)-based probe weighting, and the full MONTE model incorporating top-N probe selection. Across cancer types, each successive component resulted in significant improvements in purity estimation accuracy as measured by Pearson correlation (Figure 2A, one-sided Wilcoxon signed-rank test; MONTE vs OLS p=9.5e-7, MONTE vs Var p=1.4e-6, MONTE vs SNR p=4.5e-3). Although the magnitude of improvement varies across cancer types, MONTE consistently outperforms or matches simpler variants with no clear dependence on cohort size. Larger gains are observed primarily in cancers with lower baseline OLS performance, suggesting that MONTE preferentially stabilizes purity estimation in more challenging settings.

**Figure 2.**
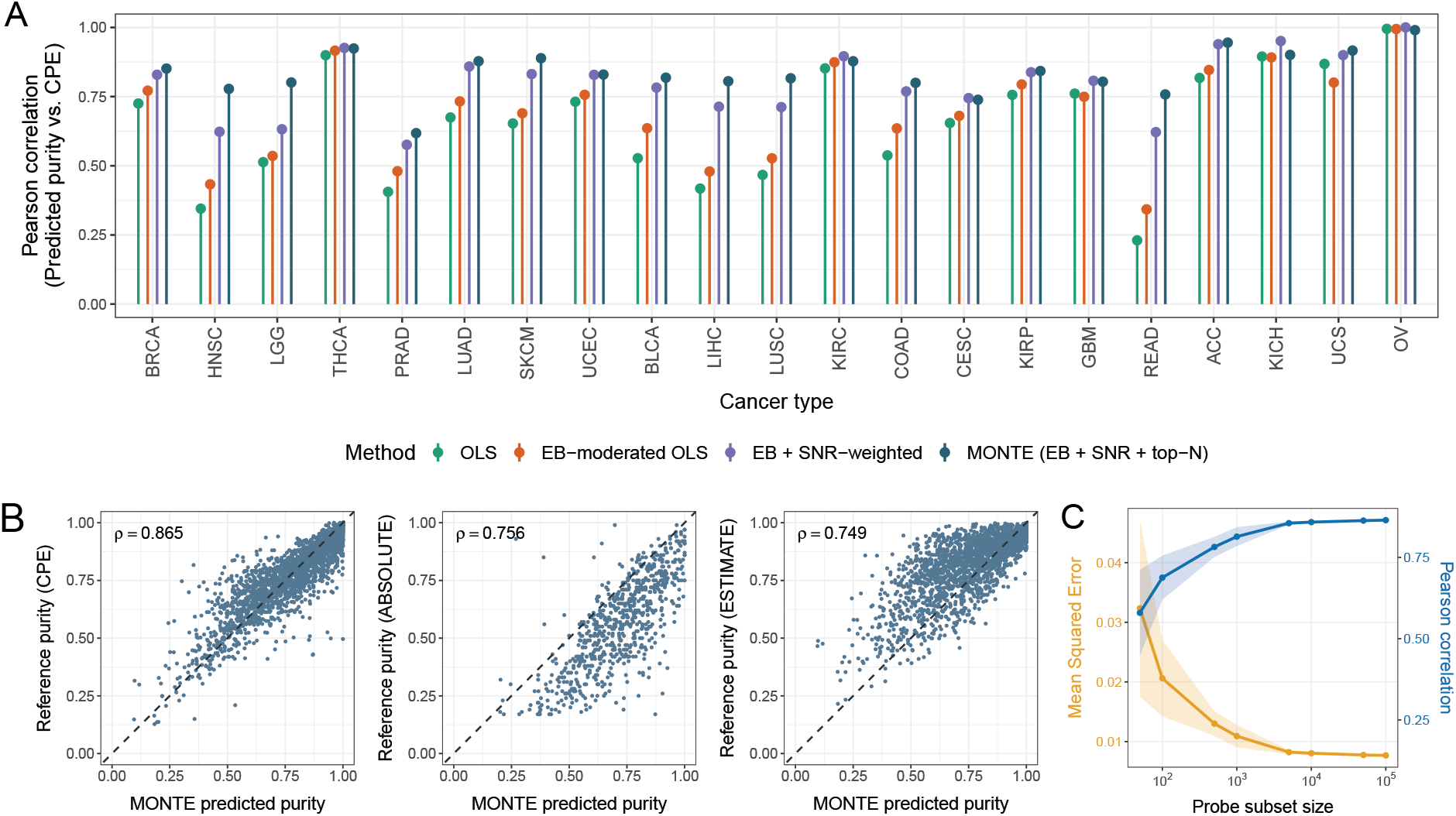
A single pan-cancer MONTE model accurately estimates tumor purity across cancer types. **(A)** Pearson correlation between estimated and reference tumor purity (CPE) on a held-out TCGA test set, shown across cancer types for 4 model variants: baseline ordinary least squares (OLS), OLS with empirical Bayes variance moderation (EB-moderated OLS), EB with signal-to-noise weighted probe aggregation (EB + SNR-weighted), and the full MONTE model with top-N probe selection (EB + SNR + top-N). The top_n hyperparameter was selected through 5-fold cross-validation on the training set, resulting in optimal top_n of 250. Each successive component improves estimation performance. **(B)** Scatter plots of MONTE predictions on the TCGA test set against CPE (left), ABSOLUTE (middle), and ESTIMATE (right) reference purity values. *ρ* denotes Pearson correlation. **(C)** Pearson correlation (blue) and mean squared error (yellow) between predicted and reference purity as a function of the number of probes sampled from the input. Shaded regions indicate standard deviation across five random seeds. Performance stabilizes at around 1000 probes.

The full MONTE model achieves strong agreement with reference purity values on the held-out test set (Figure 2B; Pearson *ρ*=0.865, MSE=0.008), demonstrating that a single pan-cancer model can accurately estimate tumor purity without using cancer labels. To assess whether this performance is driven by genuine purity-associated methylation signal rather than recapitulation of the methylation-based component of CPE, we further compared MONTE predictions for the held-out test samples against the ABSOLUTE and ESTIMATE reference purity measures on the subset of samples for which each was available. MONTE predictions agree closely with both references (Figure 2B; Pearson *ρ*=0.756, n=729 against ABSOLUTE; *ρ*=0.749, n=1,972 against ESTIMATE), exceeding the agreement between LUMP and each of these references on the same samples (Supplementary Figure 1, Pearson *ρ*=0.649 against ABSOLUTE and Pearson *ρ*=0.638 against ESTIMATE). We also examined whether MONTE’s top ranked probes simply recapitulate LUMP-associated methylation signal. LUMP uses 44 predefined probes to calculate its reference purity measure, of which only 11 probes were present in our training dataset after quality control. Among MONTE’s top ranked probes (n=250), only 1 overlapped with the LUMP probe set, indicating that MONTE identifies a broader and largely distinct set of purity-associated probes across the pan-cancer cohort.

To assess robustness to probe availability, we further evaluated MONTE using randomly subsampled input probe sets across several orders of magnitude, with ten random seeds per setting. Performance improves rapidly as the number of probes increases and stabilizes within a few thousand probes (Figure 2C), corresponding to <1% of the ∼163K filtered probes captured on the array, showing that MONTE is robust to substantial probe filtering and does not rely on predefined probe panels.

### MONTE improves the generalizability and scalability of pan-cancer tumor purity estimation

We next compared MONTE with existing DNA methylation–based tumor purity estimation methods to assess applicability, accuracy, and computational scalability in a pan-cancer setting (Figure 3). Because existing methods rely on cancer-specific reference models or probe sets, we evaluated each under two settings: pretrained and dataset-fitted. In the pretrained setting, methods were applied using the provided reference models or probe sets, reflecting their intended out-of-the-box usage, though we note that all of these methods use TCGA data in their training and therefore may inevitably overlap with our test samples.

**Figure 3.**
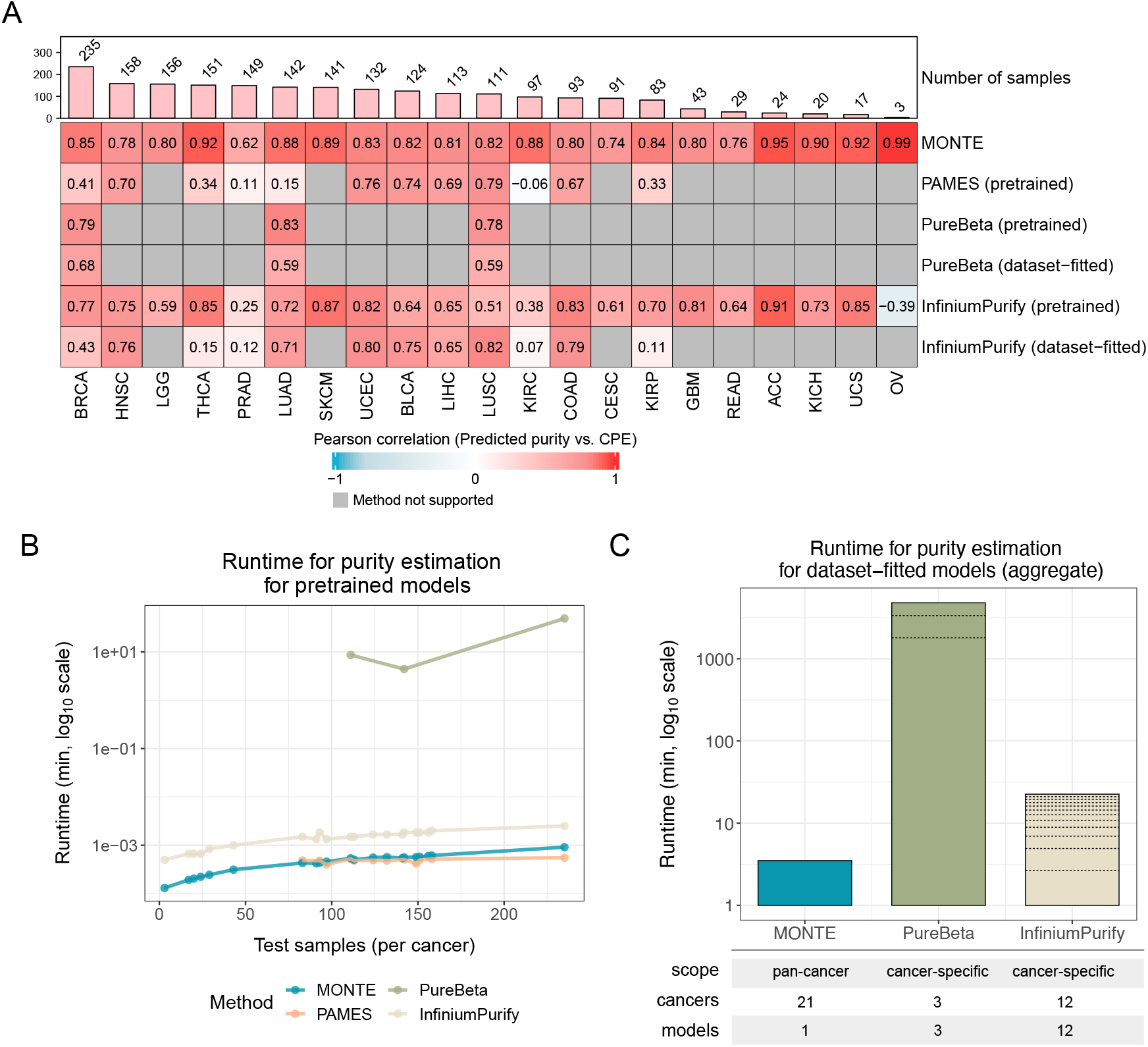
MONTE outperforms cancer-specific methods at substantially lower computational cost. **(A)** Pearson correlation between predicted purity and CPE on the TCGA test set, by cancer type, for MONTE and three existing methods (PAMES, PureBeta, InfiniumPurify) under pretrained and dataset-fitted evaluation settings. Pretrained models use the published reference models or differentially methylated CpG sets provided with each method, which may include samples used in our test set in their original training; dataset-fitted models are trained on the same TCGA train/test split used for MONTE. Sample counts per cancer type are shown above. Gray cells indicate cancer types not supported by the given method. **(B)** Runtime using pretrained models for per-cancer purity estimation plotted as a function of the number of test samples per cancer type. **(C)** Total runtime for dataset-fitted purity estimation across all 21 cancer types for MONTE, 12 cancer types for InfiniumPurify, and 3 cancer types for PureBeta in the test set for each method with fitting functionality. The dashed lines in the stacked bar represent the individual runtime for each cancer type trained, and the full runtime table is in Supplementary Table 2.

In the dataset-fitted setting, methods were retrained using the same TCGA training split used for MONTE, providing a more appropriate comparison under matched data availability. All methods were evaluated on the same TCGA test samples (held out from MONTE and the dataset-fitted settings). Coverage of competing methods is structurally limited, as pretrained PureBeta models are available for only 3 cancer types (BRCA, LUAD, LUSC), and dataset-fitted InfiniumPurify requires at least 20 tumor and 20 normal samples, restricting its appllicability in cancers with small cohorts or limited normal references.

Across cancer types, MONTE achieves consistently stronger performance than competing methods in nearly every cancer type where direct comparison is possible (Figure 3A). Specifically, MONTE significantly outperformed PAMES (difference in medians = 0.289, one-sided Wilcoxon signed-rank test *p* = 1.26 * 10^−3^), and both variants of InfiniumPurify (difference in medians = 0.162 (dataset-fitted) and 0.129 (pretrained), one-sided Wilcoxon signed-rank test *p* = 1.63 * 10^−3^ and *p* = 8.7 * 10^−5^). The performance gain relative to PureBeta was also positive (differences in medians = 0.23 (dataset-fitted) and 0.05 (pretrained)), though statistical testing was not performed due to the limited number of cancer types for which PureBeta models were available (n=3). Notably, MONTE achieves this performance using a single pan-cancer model applied uniformly across all cancer types, whereas existing approaches rely on cancer-specific reference models or probe sets that limit their generalizability. We noticed that many of MONTE’s largest gains relative to competing methods occur in cancers with smaller training cohorts, where cancer-specific approaches are most data-limited.

We further evaluated the computational scalability of MONTE relative to existing methods for purity estimation using pretrained models as well as fitting and estimating purity with a given input dataset. Using pre-trained models, PureBeta required significantly more computation, reflecting the higher cost of its mixture modeling-based fitting procedure (Figure 3B). In the dataset-fitted setting, where each method must learn from the provided training data before estimating purity, MONTE’s pan-cancer model translates into a substantial computational advantage when applied at scale, completing both fitting and prediction across all 21 cancer types in 3.1 minutes. In contrast, PureBeta and InfiniumPurify require separate fitting procedures for each cancer type, accumulating to nearly 80 hours and 22.5 minutes, respectively, both multiple orders of magnitude slower than MONTE.

### Bayesian transfer learning enables purity estimation for external cohorts with alternative purity definitions

While the pan-cancer MONTE model provides accurate tumor purity estimates against TCGA reference purity measures, Bayesian transfer learning allows the model to be further recalibrated to external cohorts with differing definitions of purity. This framework updates probe-level coefficients from a pan-cancer prior using a limited number of samples with known purity in the external cohort, enabling efficient adaptation to new datasets and alternative purity definitions without retraining the model from scratch.

To assess whether MONTE’s Bayesian transfer learning generalizes to external cohorts, we evaluated it on 3 cohorts from the Pan-Cancer Analysis of Whole Genomes (PCAWG) [15, 16], where reference purity measures were derived from whole-genome sequencing (WGS): ovarian (OV, n=70), pancreatic adenocarcinoma (PACA, n=93), and prostate adenocarcinoma (PRAD, n=98). PCAWG provides a particularly strong validation setting because WGS-derived purity is computed from the tumor genome without using methylation signals, and none of these donors were profiled in TCGA. We performed 5-fold cross-validation, where for each fold, the pan-cancer prior was recalibrated by Bayesian transfer learning using the remaining four folds and evaluated on the held-out fold. Reported correlations and mean squared errors are computed on the pooled held-out predictions across all folds.

Without recalibration, MONTE showed moderate to strong agreement with PCAWG purity estimates (Figure 4A; OV Pearson *ρ*=0.774, PACA *ρ*=0.688, PRAD *ρ*=0.563), suggesting that the pan-cancer model already captures purity-associated signal that generalizes beyond TCGA and methylation-based references.

**Figure 4.**
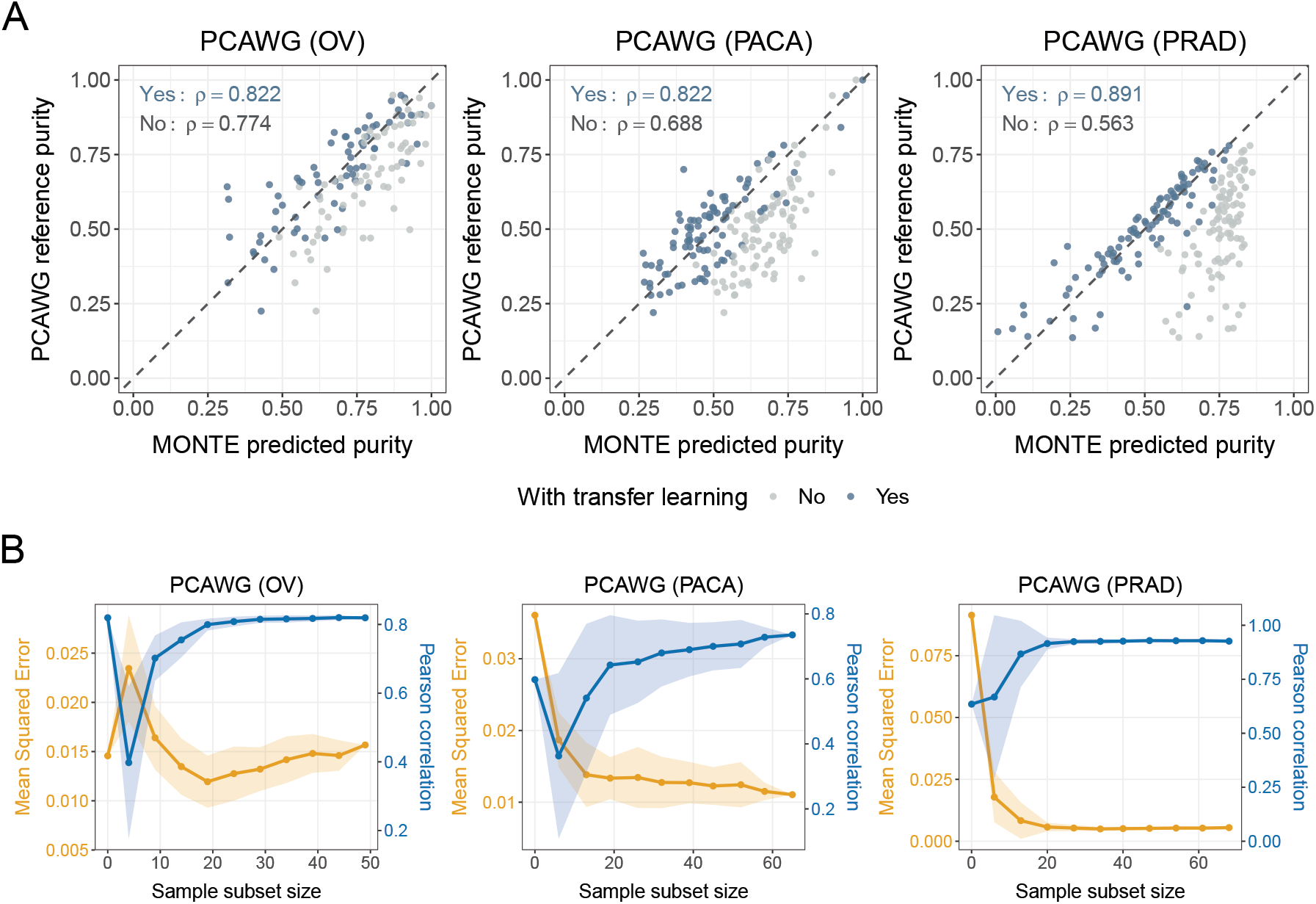
Bayesian transfer learning enables sample-efficient recalibration to external cohorts with alternative purity definitions. (A) Purity estimation performance of MONTE before (gray) and after (blue) Bayesian transfer learning on three external PCAWG cohorts: ovarian (OV), pancreatic adenocarcinoma (PACA), and prostate adenocarcinoma (PRAD), each evaluated against PCAWG’s whole genome sequencing-derived purity measures. *ρ* denotes Pearson correlation. Bayesian transfer learning recalibrates probe-level coefficients to the external purity definition and substantially improves agreement. **(B)** Pearson correlation (blue) and mean squared error (yellow) between MONTE predicted purity and PCAWG-reported reference purity as a function of the number of samples used in Bayesian transfer learning on PCAWG-OV, PCAWG-PACA, and PCAWG-PRAD. Shaded regions indicate standard deviation across 10 random seeds. Performance stabilizes with approximately 20 calibration samples.

Bayesian transfer learning substantially improved agreement in all three cohorts (OV *ρ*=0.822, PACA *ρ*=0.822, PRAD *ρ*=0.891), correcting systematic offsets while retaining information from the original model.

We further examined how recalibration performance changes as a function of the number of samples used for Bayesian transfer learning. Both Pearson correlation and mean squared error improved rapidly with sample size, stabilizing by approximately 20 calibration samples across all 3 cohorts (Figure 4B). This sample-efficient behavior reflects the advantage of the pan-cancer prior, which already captures genome-wide relationships between DNA methylation and tumor purity, allowing effective adaptation to alternative purity definitions with limited cohort-specific data.

### MONTE corrects bulk methylation profiles toward tumor-intrinsic values

The pan-cancer MONTE model and its Bayesian transfer learning extension produce probe-level coefficients that quantify the contribution of tumor purity to methylation at each CpG. These coefficients can be used to project bulk methylation measurements toward a target tumor purity, yielding CpG-resolved corrected methylation profiles (see Methods). A practical advantage of the empirical Bayes framework is that it roduces moderated test statistics and per-probe significance estimates, which allows correction to be restricted to probes with statistically supported purity associations (FDR adjusted p-values ≤ 0.05), without requiring ad hoc thresholding.

To evaluate methylation correction in a cancer-specific setting while avoiding data leakage, we examined TCGA-BRCA, LUAD, and LUSC using pan-cancer priors constructed from training data with the target cancer excluded (see Methods). These cancer types were selected because they are the only ones for which both PureBeta and InfiniumPurify can be applied for direct comparison. In BRCA, MONTE-corrected methylation values produced a tighter distribution of probe-purity correlations centered near zero compared to the broad distribution observed for uncorrected values (Figure 5A). This behavior is consistent with the expectation that once the tumor and non-tumor components are disentangled, probe-level methylation should no longer track sample-level purity. For a more granular comparison across methods, probes were stratified by their uncorrected purity correlations with CPE, and we quantified the extent to which each method reduced these correlations (Figure 5B). Across all correlation bins, MONTE consistently produced the strongest reduction toward zero, with the most pronounced differences observed for probes most strongly confounded by purity in the uncorrected data. These trends were consistent in LUAD and LUSC (Supplementary Figure 2).

**Figure 5.**
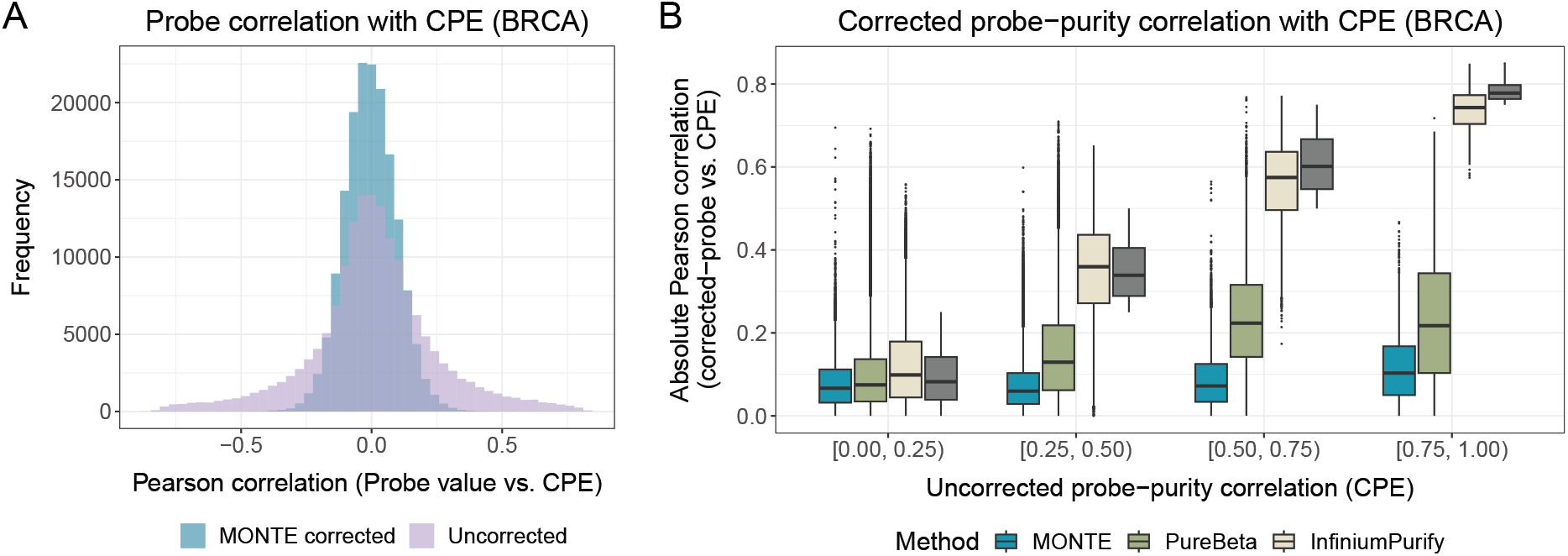
MONTE corrects bulk methylation profiles toward tumor-intrinsic signal. **(A)** Distribution of probe-purity correlations in BRCA test samples before and after MONTE correction. Correction reduces purity-associated variation, producing a tighter distribution centered near zero. **(B)** Probes are grouped by their uncorrected purity correlation with reference purity (CPE), and the resulting correlations after correction are compared for MONTE, PureBeta, and InfiniumPurify, with the uncorrected distribution shown for reference. MONTE produces the largest reduction toward zero across all correlation bins.

### Methylation correction enhances differential methylation consistency and tumor-specific signal

We examined the impact of MONTE’s methylation correction on the reproducibility of differential methylation analyses across cancer types. For each cancer, tumor samples were randomly divided into two equal halves (five random splits), and differential methylation against normal samples was computed independently for each split using both corrected and uncorrected methylation profiles. Concordance between halves was quantified as the Pearson correlation of probe-level effect sizes, and median correlations were compared before and after correction. Across nearly all cancer types, MONTE correction increased between-split concordance (Figure 6), with PRAD and LIHC being the only two exceptions with modest differences which are all less than 10^−3^. THCA showed the largest improvement (from 0.942 to 0.981), which we speculate may be due to its high immune infiltration rate [17], producing larger sample-to-sample purity variation and therefore larger gains from correction.

**Figure 6.**
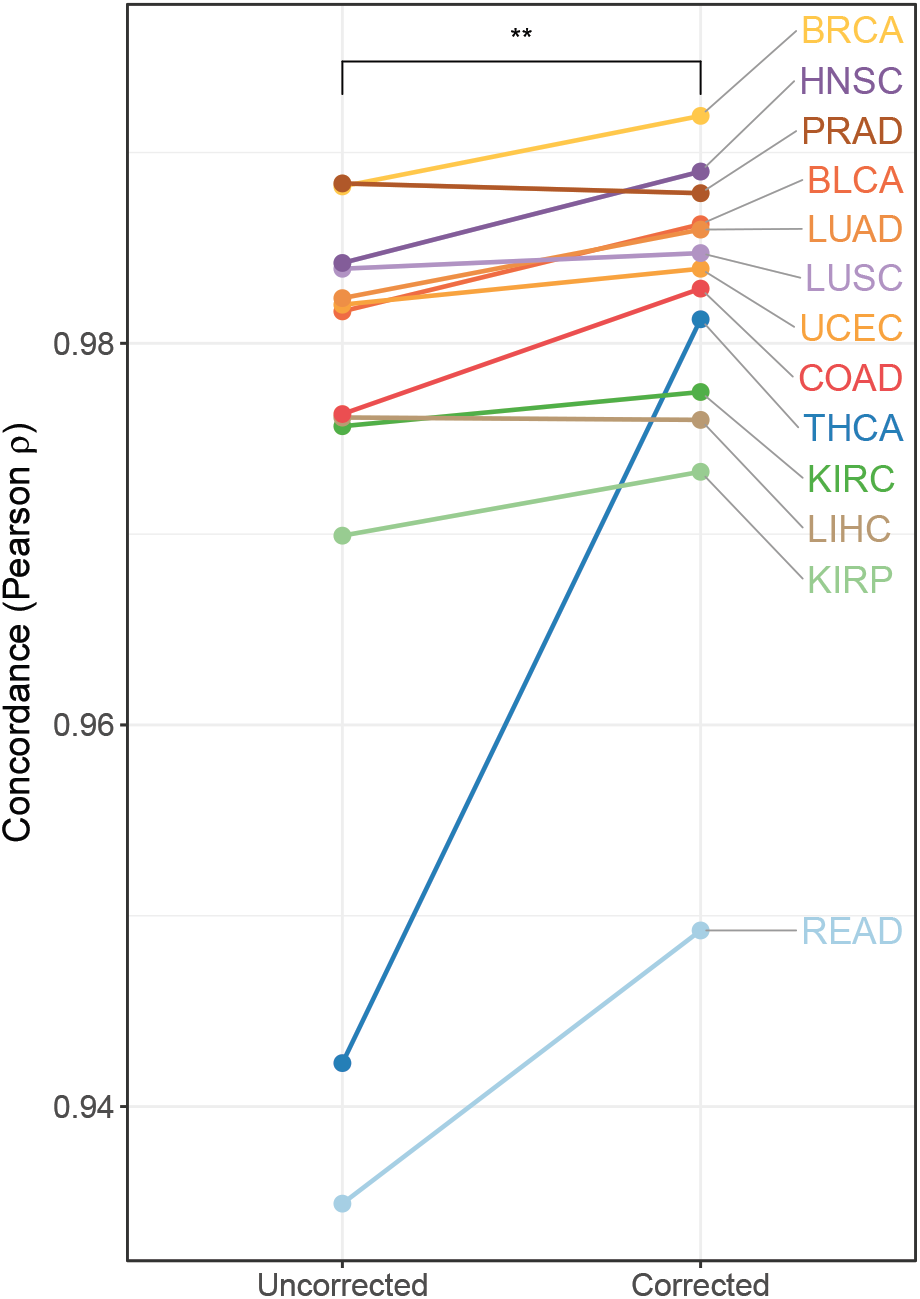
Methylation correction amplifies tumor-specific signal and improves consistency of differential methylation analyses. Median Spearman correlation between probe-level differential methylation effect sizes compared on two randomly split halves of tumor samples (across 5 random splits), shown before (Uncorrected) and after (Corrected) MONTE correction. Each line represents one cancer type; only cancer types with at least 5 tumor samples and 5 normal samples in the test set are shown (n cancers=13). Correction increases between-split concordance in nearly all cancers. (**: one sided Wilcoxon signed-rank test p-value<1e-2.)

We next asked whether MONTE correction also enhances recovery of biologically meaningful tumor-specific signal. We focused on BRCA, where well-characterized breast cancer marker genes are available, and performed differential methylation analysis between the full test set of tumor versus normal samples pre- and post-correction. For each CpG site, effect size differences were calculated as the absolute changes in differential methylation coefficients between the two settings. CpGs located in the promoter regions of literature-curated breast cancer marker genes [18] exhibited significantly larger increases in differential methylation effect sizes compared to size-matched CpG sets sampled from promoter regions of non-marker genes (permutation test with 5000 permutations, p=1.99e-4), demonstrating the amplification of tumor-relevant regulatory signal following correction.

## Discussion

In this study, we present MONTE, a unified framework that addresses several long-standing challenges in the analysis of bulk DNA methylation data by jointly enabling accurate tumor purity estimation, CpG-resolved methylation correction, and flexible adaptation across cancer types and datasets. MONTE extends classical linear modeling approaches by incorporating probe-wise empirical-Bayes variance moderation and signal-to-noise weighting, stabilizing effect estimates across heterogeneous CpGs. Compared to existing approaches, MONTE provides improved accuracy and broader applicability while remaining computationally efficient and interpretable.

A key feature of MONTE is the ability to operate as a single pan-cancer model without requiring tumor labels or normal samples, achieving strong out-of-the-box purity estimation performance across diverse cancer types. Bayesian transfer learning further extends this framework by allowing probe-purity relationships to be recalibrated using limited samples in new contexts without retraining the model from scratch. This design is particularly advantageous for underrepresented and external cohorts, where large reference datasets may not be available. Beyond purity estimation, MONTE enables CpG-resolved correction of methylation profiles, facilitating downstream analyses aimed at recovering tumor-intrinsic epigenetic signal.

As with any modeling framework, MONTE operates under a set of assumptions that define its current scope. The method models bulk methylation measurements as a linear combination of tumor and non-tumor signal, corresponding to a two-component mixture. This abstraction is intentionally simple, capturing the dominant source of confounding in bulk tumor data while remaining interpretable and data-efficient. Nevertheless, more expressive extensions, such as explicit modeling of multiple non-malignant compartments or incorporation of uncertainty in purity estimates, could further refine this framework and enable deeper understanding of tumor-microenvironment interactions.

More broadly, MONTE provides a statistical scaffold for integrating DNA methylation data. Although our analyses have focused on array-based methylation data here due to its broader availability, MONTE’s design is not platform-specific and could be extended to sequencing-based assays through the same transfer learning framework. Thus, MONTE is particularly well-suited for integrative cancer epigenomics studies across heterogeneous data to more effectively understand tumor-intrinsic epigenetic signal at scale.

## Methods

### Data preprocessing

#### Sample downloading and curation

All publicly available DNA methylation data from The Cancer Genome Atlas (TCGA) [14] were collected using the TCGAbiolinks package [19] in R. For each TCGA project, Illumina HumanMethylation450 (450K) beta values processed and normalized by the TCGA consortium were used directly. Samples without CPE purity annotations were excluded from analysis. CpG probes with missing beta values or located on sex chromosomes were excluded, following best practices for quality control [20]. For pan-cancer modeling, the intersection of probes across all projects with purity annotations was used.

#### Sample partitioning

All samples were first divided into training and test sets using a 70:30 split stratified by cancer type. Cancer type labels were used for stratification, partitioning for cancer-specific methylation correction, and per-cancer reporting of results. For pan-cancer purity estimation, the model was fit on the full training set and evaluated on the full test set, without cancer type labels ever being provided as input. For methylation correction, the pan-cancer model was additionally recalibrated to each target cancer, which required a further partition within the training set. For a given target cancer, a pan-cancer prior was constructed from training samples of all non-target cancers, and the prior was updated by Bayesian transfer learning using training samples of the target cancer, and the recalibrated model was evaluated on test samples of the target cancer (Supplementary Table 1). The topN and *τ* ^2^ hyperparameters were selected by 5-fold cross-validation on the training samples only. Because several TCGA cancers share a tissue of origin, excluding only related cancer types from prior construction would leave closely related samples that could result in leakage, so we excluded related cancer types as a group: including cancers of the digestive tract (COAD and READ), kidney (KIRC, KIRP, and KICH), uterine (UCEC and UCS), brain (GBM and LGG), and lung (LUAD and LUSC).

#### External datasets

We used data from the PRAD-CA, OV-AU, PACA-AU projects from the Pan-Cancer Analysis of Whole Genomes (PCAWG) cohort [15, 16] as an external validation, which provides 450K data with purity estimates derived from whole-genome sequencing (WGS) from cohorts independent of TCGA (the selected studies were not based in the US). Only donors with a single tumor WGS sample was used to ensure one-to-one correspondence between methylation and purity measurements.

### Model design of MONTE

MONTE models probe-wise associations between bulk DNA methylation and tumor purity using a linear regression framework. Let *X* ∈ ℝ^*n×m*^ denote centered methylation measurements (can be beta or M-values) for *n* samples and *m* CpG probes, and let *p* ∈ ℝ^*n*^ denote the vector of centered tumor purity values. For each probe *j*, MONTE fits a linear model:

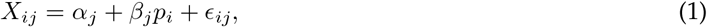

where *α*_*j*_, *β*_*j*_ are probe-specific intercept and coefficient parameters, and *ε*_*j*_ ∼ *N* (0, *σ*_*j*_) is the probe residual error and the *σ*_*j*_ is population variance for the probe *j*. Probe-wise coefficients are estimated using ordinary least squares (OLS), treating tumor purity as the independent variable and methylation values as the response.

To stabilize estimates across heterogeneous probes and variable sample sizes, MONTE applies empirical Bayes variance moderation to the OLS residual variances, shrinking probe-specific variance estimates toward a shared prior. As in limma [21], we assume that the population variance of each probe, *σ*_*j*_, is drawn from a prior distribution, an inverse chi-squared distribution:

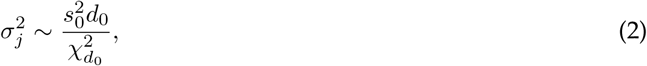

where *s*_0_ is the pooled variance estimate and *d*_0_ is the prior degrees of freedom.

The expectation of the probe’s variance *σ*_*j*_ given the observed variance *s*_*j*_ is

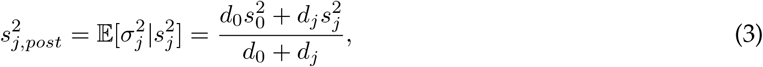

where *d*_*j*_ is the degrees of freedom for the probe. Using the posterior variance 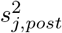, we can obtain the moderated t-statistics for each probe

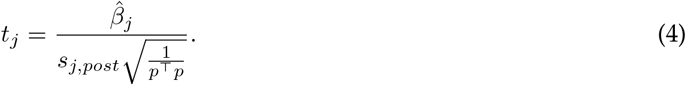

### Purity estimation with signal-to-noise (SNR) weighting

Given new samples with methylation measurements *X*^*′*^ ∈ ℝ^*n′×m′*^, where *n*^*′*^ is the number of samples and *m*^*′*^ is the number of features, MONTE estimates tumor purity using the intersection of probes between the trained model and the input data, allowing pre-trained models to be appleid across datasets with partially overlapping probe sets. After centering *X*^*′*^ using training set means, sample-wise purity estimates can be solved using a weighted least squares objective function

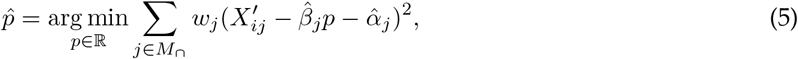

where *M*_∩_ denotes the intersecting probe set.

MONTE uses signal-to-noise (SNR) [22] based weights

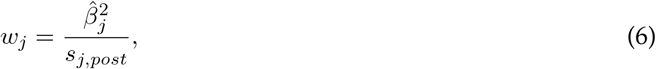

which upweight probes with strong purity association and low residual variance while downweighting noisy or weakly associated probes. This yields the closed-form purity estimator

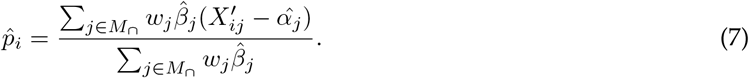

To reduce the influence of probes weakly associated with purity, MONTE optionally restricts purity estimation to the top *n* probes ranked by their moderated t-statistics. The number of probes can be specified by the user or selected via a built-in cross validation function. This built-in function automatically runs k-fold cross validation (default: k=5) on the training set across a set of *n* values (default: 5, 10, 25, 50, 100, 250, 500, 1000, 2500, 5000, 10000, 25000, 50000, 100000, 150000) and returns the *n* that corresponds to the lowest average mean squared error across folds.

### Bayesian transfer learning

MONTE supports adaptation to new datasets through Bayesian transfer learning, which updates probe-level purity coefficients using limited samples with known purity. We assume a Gaussian distribution on 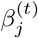 for probe *j* in the new dataset, 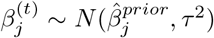, where *τ*^2^ captures the uncertainty of the pan-cancer coefficient estimate for probe *j*.

Given a target dataset with labeled samples, probe-level coefficients 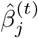 and associated variance 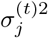 are estimated using the same linear model. The posterior distribution of 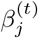 thus has the closed-form mean

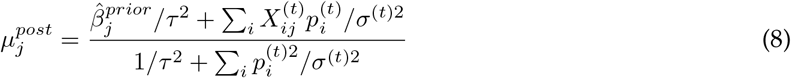

and variance,

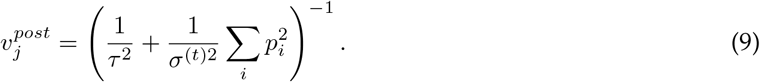

This update corresponds to a precision-weighted combination of the pan-cancer prior and the target-specific estimate, allowing probe-purity relationships to be recalibrated efficiently without retraining the model from scratch.

### Correction of methylation values

MONTE enables CpG-resolved correction of bulk DNA methylation profiles by correcting observed methylation values to a target tumor purity. Given estimated sample purity 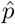 and probe-level coefficients 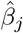, methylation values for sample *i* and probe *j* are corrected as

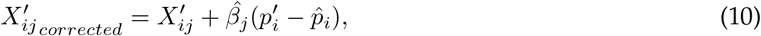

where *p*^*′*^ denotes the target purity, typically set to 1 to obtain a pure tumor methylation profile. This correction corresponds to projecting each probe’s methylation value along its estimated purity-associated direction to the desired purity level. The operation is applied efficiently in matrix form across all probes and samples. Because not all probes are necessarily strongly associated with tumor purity, MONTE provides optional controls to restrict correction to informative probes based on probe-level significance thresholds. MONTE also provides easy access to detailed probe statistics that include confidence intervals, allowing users to assess the reliability of individual corrections.

### Evaluation of CpG-resolved methylation correction

To evaluate methylation correction of the pan-cancer model when cast to a specific cancer type, we trained separate pan-cancer models excluding samples from each target cancer type, followed by transfer learning on training samples of the target cancer type. Because LUAD and LUSC are both lung-related cancer types and share similar biological characteristics, both cancer types were excluded during pan-cancer model training when either LUAD or LUSC was used as the target cancer type. Methylation correction performance was evaluated on a completely held-out test set of samples, where the previously described methylation correction procedure was applied to probes with an FDR-adjusted p-value ≤ 0.05. We repeat this procedure for all 21 TCGA cancer types and release all models, but we focus evaluations on BRCA, LUAD, and LUSC because they are the only ones that both PureBeta and InfiniumPurify are able to be used.

### Differential methylation, effect size, and subset concordance

Differential methylation (DM) was performed using the *methylize* [23] package in Python (v3.13.3). Significant probes were defined using FDR-adjusted p-values (*<* 0.05). CpGs were mapped to promoter regions (±1.5 kb from transcription start sites) using the hg38 manifest [20].

For each CpG probe, tumor–normal effect sizes were obtained from the linear DM coefficient. Absolute effect size changes were calculated as the difference between corrected and uncorrected coefficients. Cancer marker genes were obtained from MethMarkerDB [18]. Marker-associated CpGs were defined as probes mapped to promoters of curated marker genes. To assess whether marker CpGs exhibited larger effect size increases than compared to those of random non-marker sets, a permutation test was performed by randomly sampling non-marker probes 5,000 iterations.

To assess the consistency of DM results between corrected and uncorrected methylation, test tumor samples were randomly split into two halves five times for each cancer type. For each split, DM was performed against normal samples using *methylize* [23]. Only cancer types with at least five tumor and five normal samples were included. Concordance between splits was quantified using Spearman correlation of effect sizes. Median correlation across iterations was compared between corrected and uncorrected data.

### Existing purity estimation and correction methods

We compared MONTE against three existing DNA methylation array–based tumor purity estimation tools and their corresponding beta value correction procedures: PAMES [7], InfiniumPurify [5], and PureBeta [6]. All methods were run using pretrained and dataset-fitting modes following recommended settings whenever possible. We note that all methods include TCGA data in their training, so the pretrained models inevitably have the advantage of being trained on some of the testing samples.

PAMES was evaluated using the provided informative probe collections, which supports 12 cancer types (BLCA, BRCA, COAD, HNSC, KIRC, KIRP, LIHC, LUAD, LUSC, PRAD, THCA, UCEC). PureBeta under both pretrained and dataset-fitted settings. Pretrained models used the precomputed regression parameters provided for BRCA, LUAD, and LUSC. Dataset-fitting was also performed for these same cancer types, which the authors have described as satisfying the method’s modeling assumptions. For InfiniumPurify, pretrained analyses were restricted to tumor types supported by the package so READ was missing, but as suggested in the method documentation, we used the COAD model for READ predictions. In the dataset-fitting setting, model fitting required at least 20 tumor and 20 normal samples, which limited applicability in rarer cancer types with small sample sizes. Beta value correction also required at least 20 normal samples, restricting the number of cancer types that we could compare against.

We additionally attempted to evaluate RF_purify [9] and MEpurity [8]; however, RF_purify could not handle missing values introduced during standard quality control, and MEpurity produced empty outputs when run according to the provided documentation. These methods were therefore excluded from further analysis.

## Data and code availability

Our implementation of MONTE package is available on GitHub at https://github.com/ylaboratory/MONTE, with additional analysis code, tutorials, and all TCGA models available at https://github.com/ylaboratory/MONTE-analysis, all released under the BSD 3-clause license for open source use.

## Declaration of interests

The authors declare no competing interests.

## Acknowledgments

The authors would like to thank members of the ylaboratory for helpful discussions. This work was supported by the Cancer Prevention & Research Institute of Texas [CPRIT RR190065 to VY] and the National Science Foundation [NSF DBI-2144534 to VY]. VY is a CPRIT Scholar in Cancer Research.

## Notes

### Competing Interest Statement

The authors have declared no competing interest.

### Summary of Updates

Updated the manuscript title; Improved the figure visualization; Improved the writing. Added a supplementary table 2.

