## Supplementary Material for "MONTE enables unified pan-cancer tumor purity estimation and methylation correction from bulk DNA methylation arrays"

### MONTE: Supplementary Materials

#### Supplementary Tables

| Cancer | $N_{train}$ | $N_{transfer}$ | $N_{test}$ | Best topN | Best $\tau^2$ |
|---|---|---|---|---|---|
| BRCA | 2501 | 314 | 235 | 250 | 100.00 |
| HNSC | 2604 | 211 | 158 | 500 | 100.00 |
| LGG | 2550 | 208 | 156 | 500 | 10.00 |
| THCA | 2614 | 201 | 151 | 250 | 100.00 |
| PRAD | 2616 | 199 | 149 | 250 | 0.01 |
| LUAD | 2478 | 189 | 142 | 250 | 0.10 |
| SKCM | 2628 | 187 | 141 | 500 | 100.00 |
| UCEC | 2617 | 175 | 132 | 250 | 100.00 |
| BLCA | 2649 | 166 | 124 | 250 | 100.00 |
| LIHC | 2665 | 150 | 113 | 100 | 0.01 |
| LUSC | 2478 | 148 | 111 | 250 | 100.00 |
| KIRC | 2549 | 130 | 97 | 250 | 100.00 |
| COAD | 2652 | 123 | 93 | 250 | 100.00 |
| CESC | 2693 | 122 | 91 | 250 | 10.00 |
| KIRP | 2549 | 110 | 83 | 250 | 100.00 |
| GBM | 2550 | 57 | 43 | 500 | 10.00 |
| READ | 2652 | 40 | 29 | 250 | 100.00 |
| ACC | 2783 | 32 | 24 | 250 | 10.00 |
| KICH | 2549 | 26 | 20 | 250 | 1.00 |
| UCS | 2617 | 23 | 17 | 250 | 0.01 |

**Supplementary Table 1:** Sample distribution for cancer-specific MONTE with cross-validated hyperparameters. The training set contains all but the target cancer from the training split, the transfer learning set contains only the target cancer from the training split, and the test set contains only the target cancer from the test split. To avoid data leakage due to similar cancer types, we exclude cancer types from the same tissue during pan-cancer model training (see Methods for more details).

| Method | Scope | Cancer | Runtime (min) |
|---|---|---|---|
| MONTE | Pan-cancer | All | 3.50 |
| PureBeta | Cancer-specific | BRCA | 1801.17 |
| | Cancer-specific | LUAD | 1554.38 |
| | Cancer-specific | LUSC | 1442.37 |
| InfiniumPurify | Cancer-specific | BRCA | 2.66 |
| | Cancer-specific | KIRC | 2.25 |
| | Cancer-specific | HNSC | 2.04 |
| | Cancer-specific | THCA | 1.96 |
| | Cancer-specific | PRAD | 1.93 |
| | Cancer-specific | UCEC | 1.81 |
| | Cancer-specific | LUAD | 1.77 |
| | Cancer-specific | LIHC | 1.71 |
| | Cancer-specific | LUSC | 1.66 |
| | Cancer-specific | BLCA | 1.62 |
| | Cancer-specific | COAD | 1.57 |
| | Cancer-specific | KIRP | 1.53 |

**Supplementary Table 2:** Training runtime for dataset-fitted tumor purity estimation methods. MONTE was trained once using the pan-cancer dataset, whereas PureBeta and InfiniumPurify were trained separately for each available cancer type. Runtime is reported in minutes for each individual model fit.

#### Supplementary Figures

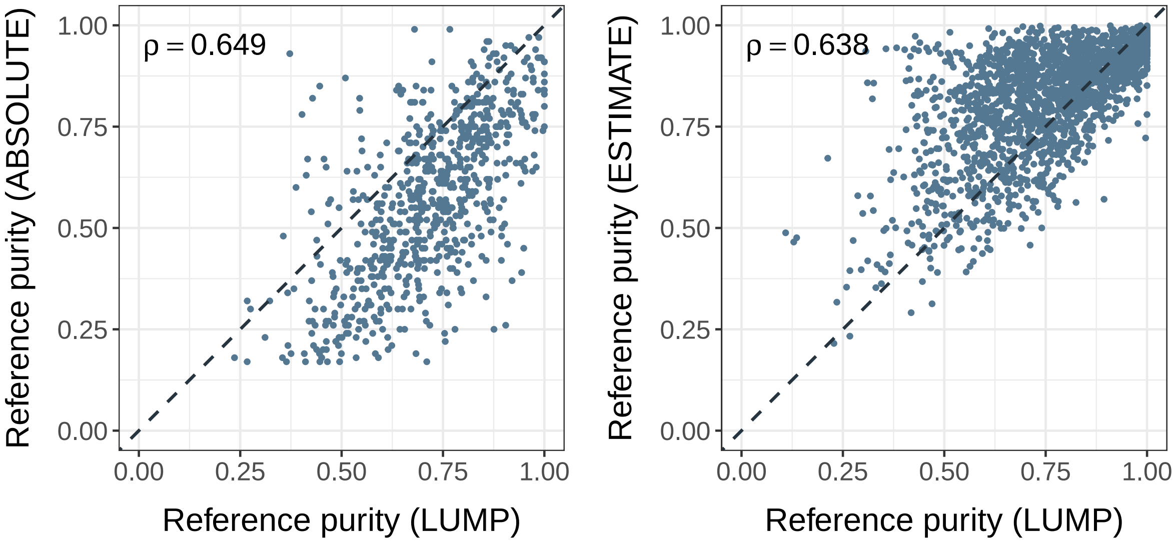


**Supplementary Figure 1: Comparison of correlations with different metrics against LUMP in TCGA test set.** Scatterplots of LMUP reference purity values on the TCGA test set against ABSOLUTE (left), and ESTIMATE (right) reference purity values. $\rho$ denotes Pearson correlation.

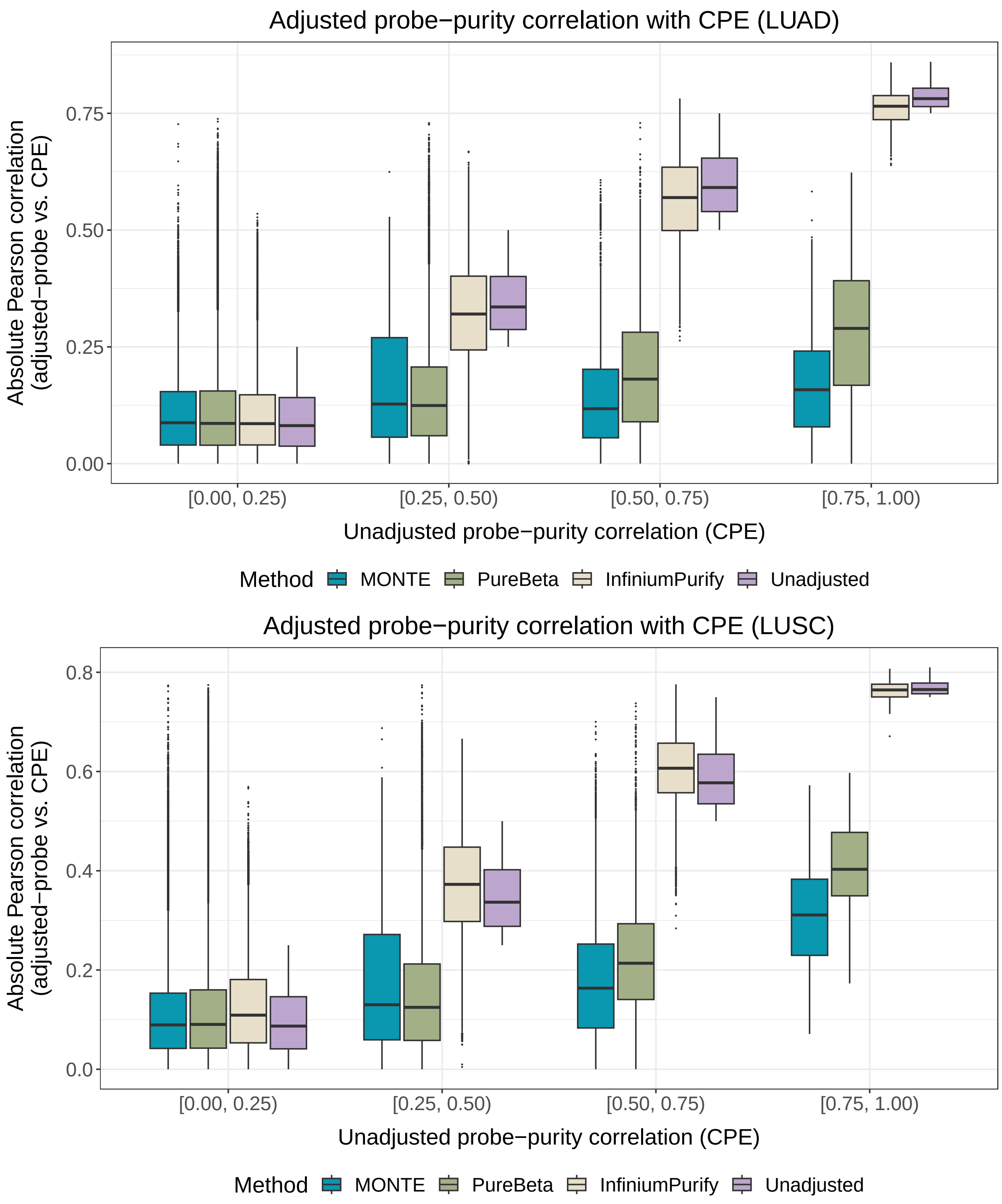


**Supplementary Figure 2: Probes grouped by purity before and after purification.** Probes grouped by correlation with purity before purification with respective purified correlations for each benchmarking method. These cancers are shown because they represent the subset in which both PureBeta and InfiniumPurify can be applied.
